# Mixed-Feedback Architectures for Precise Event Timing Through Stochastic Accumulation of Biomolecules

**DOI:** 10.1101/2023.05.22.541681

**Authors:** Sayeh Rezaee, César Nieto, Abhyudai Singh

## Abstract

The timing of biochemical events is often determined by the accumulation of a protein or chemical species to a critical threshold level. In a stochastic model, we define event timing as the first-passage time for the level to cross the threshold from zero or random initial conditions. This first-passage time can be modulated by implementing feedback in synthesis, that is, making the production rate an arbitrary function of the current species level. We aim to find the optimal feedback strategy that reduces the timing noise around a given mean first-passage time. Previous results have shown that while a no-feedback strategy (i.e., an independent constant production rate) is optimal in the absence of degradation and zero-molecules initial condition, a negative feedback is optimal when the process starts at random initial conditions. We show that when the species can be degraded and the synthesis rates are set to depend linearly on the number of molecules, a positive feedback strategy (the production rate increases with the level of the molecule) minimizes timing noise. However, if no constraints on the feedback are imposed, the optimal strategy involves a mixed feedback approach, which consists of an initial positive feedback followed by a sharp negative feedback (the production rate decreases with the level) near the threshold. Finally, we quantify the fundamental limits of timing noise reduction with and without feedback control when time-keeping species are subject to degradation.

## I. Introduction

Precision in the timing of biochemical events within cells is needed for high fidelity functioning despite inherently noisy processes occurring with low-copy number components. Several biological processes critically depend on timing mechanisms based on threshold-crossing of given variables. These include, neuronal firing of action potential, cell-cycle regulation, tissue development, biological clocks, apoptosis, signal transduction, and gene activation [1]–[12]. This time variability can be reduced by using self-regulation in protein synthesis rates [13]–[17]. Self-regulation strategies, also known as feedback, involve adjusting the transcription/translation rate based on the current protein level. Finding an optimal feedback technique that minimizes this timing variability in particular systems is crucial to understanding the mechanisms of cell timing, and uncovering the fundamental limits of timing noise suppression.

One of the most used approaches for modeling cell timing is based on first-passage time (FPT) statistics [18]–[25]. Different feedback regulation schemes have been explored, and different parameters have been optimized to minimize the FPT variability while keeping the mean FPT constant [13]–[17], [26]. The solution for simple systems such as constant protein production without degradation or dilution has already been studied. Interestingly, for this system, the minimum timing noise is achieved if there is no feedback [27]. Recently, we have shown that if the protein is continuously diluted, the effect of dilution can be compensated by a feedback strategy that increases the transcription rate as the protein level increases [28]. However, this model has some limitations. First, it assumes that dilution is a continuous deterministic process, but for some biochemical systems, stochastic protein degradation may be more relevant than dilution. Second, it assumes that the feedback strategy is linearly dependent on the protein level. In a general approach, the optimal feedback strategy is expected not to grow linearly with the protein amount but to be an arbitrary function of this number. To date, the general solution to this problem of optimization characterizing timing accuracy is unknown.

In this study, we address the limitations mentioned above. Specifically, we employ a modeling framework wherein protein levels are represented as discrete random variables. Moreover, protein production and degradation mechanisms are characterized as stochastic birth-death processes. The occurrence rates of these processes depend on the protein level, such that the degradation probability of a protein molecule increases linearly with the protein copy number, and the synthesis rate is an arbitrary function of this number. Our primary objective is to investigate the feedback function that minimizes noise in FPT around a predetermined fixed mean.

The article consists of the following sections: First, we review the problem and the solution in the absence of degradation with both zero-molecule initial conditions and a random initial protein number. Second, we consider optimizing the slope of a linear synthesis rate with stochastic degradation. We use the small-noise approximation to obtain analytic formulas that are valid for high threshold numbers. Third, we present a solution for the low-number regime. We analytically solve the optimization problem for two molecules and numerically solve the master equation of the system for five molecules, optimizing over the synthesis rates to minimize FPT fluctuations. As a main result, mixed feedback emerges as optimal, wherein the synthesis rate increases non monotonically with the protein level and sharply declines near the threshold. Finally, we discuss the implications of our findings for biological systems and suggest potential applications.

## II. Formulating event timing as a first-passage time problem

Let *x*(*t*) ∈ {0, 1, 2, …} be an integer-valued process representing the molecular count of a certain biochemical species at time t. The time evolution of *x*(*t*) can be modeled as a birth-death process with a constant synthesis rate *k* and a degradation rate γ. More specifically, synthesis events occur according to a Poisson process with rate *k*. In this case, the probability of copy numbers increasing from *i* to *i* + 1 in an infinitesimal time interval (*t, t* + *dt*] is given by:

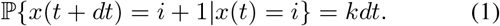

The timespan each molecule lives is an independent and identically distributed random variable that is exponentially distributed with mean timespan 1/γ, and corresponds to the following probability of degradation:

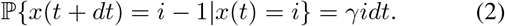

A feedback loop is implemented by allowing the production rate *k*_*i*_, to vary arbitrarily with the current levels of the species, denoted as *x*(*t*) = *i*. This alters the synthesis probability (1) to

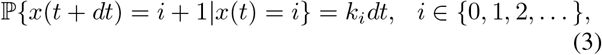

where the time required for transition from *i* to *i* + 1 molecules is exponentially distributed with mean 1/*k*_*i*_. This formalism enables the realization of various feedback scenarios:

- A constant synthesis rate *k*_*i*_ = *k* corresponds to a no-feedback strategy.
- Negative feedback corresponds to *k*_*i*_ decreasing with increasing *i*, i.e., the synthesis rate declines with higher molecular counts.
- Similarly, positive feedback corresponds to *k*_*i*_ increasing with *i*.
- Nonmonotonic shapes of *k*_*i*_ capture complex feedback strategies that combine both negative and positive feedbacks.

Let a positive integer *X* be a critical threshold level for molecular counts that triggers an event. Then, event timing is a random variable defined by the first-passage time (FPT)

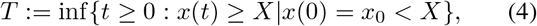

which corresponds to the first time the random process *x*(*t*) reaches *X* starting from an initial condition *x*_0_ at time *t* = 0. For a fixed *X* and degradation rate *γ*, our goal is to determine the optimal feedback strategy *k*_*i*_, *i* ∈ {0, 1, 2, … *X*−1} for synthesizing molecules that minimizes the random fluctuations in *T, while ensuring a given mean first-passage time* (Fig. 1B).

**Fig. 1:**
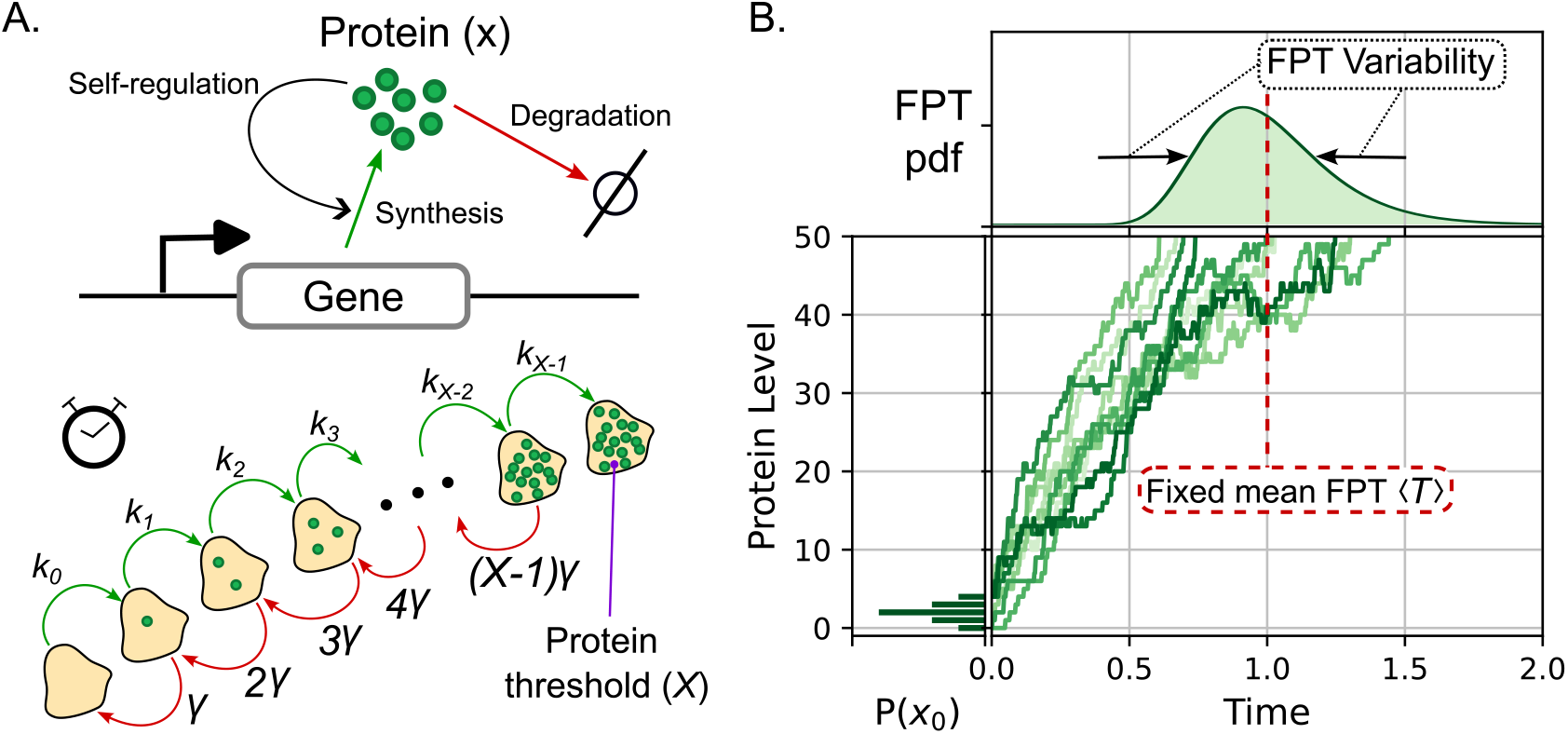
Event timing of biochemical processes triggered by the accumulation of proteins to a threshold. **A**. Model of protein synthesis and degradation from gene expression. Feedback is implemented by allowing the synthesis rate *k*_*i*_ to vary as a function of the current protein level. The stochastic dynamics of the protein count is modeled as a birth-death process. Synthesis, wherein the number of molecules jumps from *i* to *i* + 1, occurs at a rate *k*_*i*_. Additionally, the degradation, which is the jump from *i* to *i* − 1, occurs at a rate *γi*. **B**. Example trajectories of protein copy number over time starting from a random initial condition. The event is triggered when the level reaches a critical threshold. Our goal is to obtain the strategy of *k*_*i*_ that minimizes the stochastic variability in the FPT. In the optimization, we fix the threshold *X*, the degradation rate *γ*, and the average time ⟨*T*⟩ required to reach the threshold.

## III. Optimal feedback strategy in the absence of degradation

Having formulated the problem, we first focus on the special case of no degradation i.e., *γ* = 0, where molecular counts only build up over time.

### A. FPT problem with Zero-molecules initial condition

If the system starts with zero molecules with probability one at *t* = 0, the first-passage time is essentially a sum of each reaction time, which is an exponentially distributed random variable with mean 1/*k*_*i*_, *i* ∈ { 0, 1, …, X − 1}. Using the angular brackets ⟨⟩ to denote the expected-value operation, the moments of *T* follow

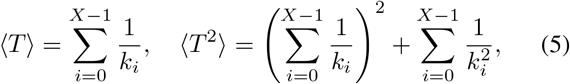

which leads to the variance as below.

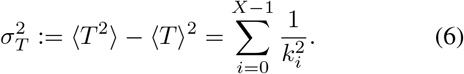

Our goal is to find the optimal sequence *k*_*i*_, *i* ∈ {0, 1, …, *X* − 1} that minimizes the variance 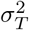 while keeping ⟨*T*⟩ fixed. To obtain the solution, we propose that optimal rates can be written as

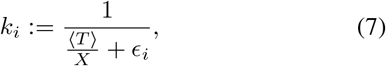

with *ϵ*_*i*_ ∈ ℝ having arbitrary values. If we fix ⟨*T*⟩ from (5), the following constraint

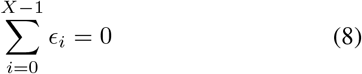

is required to ensure a fixed mean FPT. On the other hand, using (6) and (8), the variance in FPT is given by

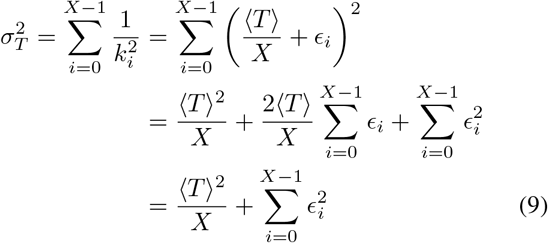

The sum of squares 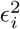 on the right-hand-side of (9) is minimized only when all *ϵ*_*i*_ = 0. Therefore, the optimal feedback strategy is to set a constant synthesis rate as:

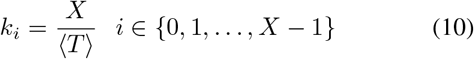

and the corresponding optimal noise in FPT

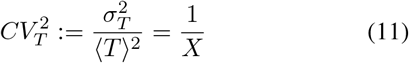

as quantified by its coefficient of variation.

In summary, when *γ* = 0, to reach a threshold at a prescribed time, *the optimal strategy is to have a constant synthesis rate, and any form of feedback will amplify timing fluctuations* [27]. Moreover, 1/*X*, the inverse of the threshold level, is the fundamental limit to which timing noise can be suppressed. Previous studies have extended this problem to consider production occurring in molecular bursts, as seen in gene expression measurement in single cells [29]–[36], that is, each synthesis event produces several molecules in contrast to just one molecule as considered here. For this bursty-birth process, the optimal feedback in the absence of degradation is similar to a no-feedback strategy, where all *k*_*i*_, i ∈ { 1, …, *X* − 1} are exactly the same except for the first synthesis rate *k*_0_ that is lower than the rest [27], [37], [38].

### B. Random initial condition

We next extend the mentioned results to consider the process starting with a random initial condition as follows.

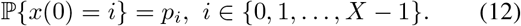

This assumption leads to the following mean FPT,

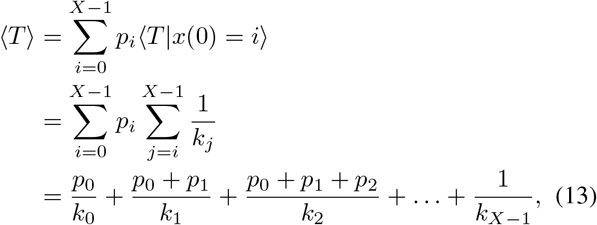

and its corresponding (uncentered) second-order moment.

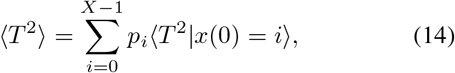

where

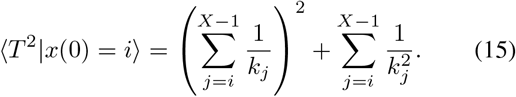

To perform the optimization, one can find k_0_ as a function of *k*_1_, …, *k*_*X*−1_ from (13) to have a fixed ⟨*T*⟩. Substituting this value of *k*_0_ into (14), one can then optimize over *k*_1_, …, *k*_*X*−1_ to minimize ⟨*T*^2^⟩. We did the optimization by simultaneously solving the equations

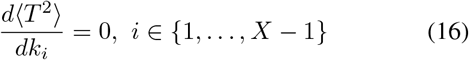

to find the rates *k*_1_, …, *k*_*X*−1_.

We illustrate the results for a threshold of *X* = 6 molecules. If the initial condition is zero molecules with probability *p*_0_ and one molecule with probability 1 − *p*_0_ then the optimal rates are

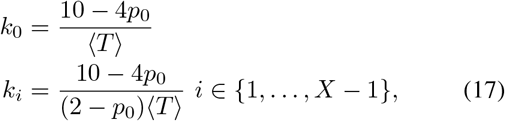

where all *k*_*i*_, *i* ∈ {1, …, *X* − 1} are equal and *k*_0_ set higher than the rest. As anticipated, when *p*_0_ →1, the result converges to having all rates as 6/ ⟨*T*⟩. In addition, if we consider an initial condition with equal probability in all states, meaning *p*_*i*_ = 1/6, *i* ∈ {0, 1, …, *X* −1}, then the optimal rates are

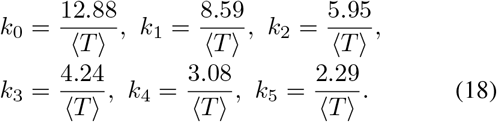

which corresponds to a negative feedback strategy with rates decreasing as *i* increases.

In the next section, we describe the optimization problem in the presence of protein degradation assuming a linear synthesis rate and a low-noise limit with a large threshold *X*.

## IV. Optimal rates for linear self-regulation at low-noise regime

In this section, we consider the accumulation of biomolecular materials starting from *x*(0) = 0 with probability 1 and the degradation of molecules with rate *γ*. To simplify the feedback strategy, we assume a linear synthesis rate *k*_*i*_ = *k*_0_ +*αi*. Here, *k*_0_ is the basal rate of protein production, and the slope *α* is called the feedback strength. In this case, positive feedback is implemented when *α* > 0, and negative feedback occurs when *α* < 0.

As a consequence of this linear stochastic formulation, the time evolution of the statistical moments of *x*(*t*) can be obtained precisely by using the moment dynamics formalism.

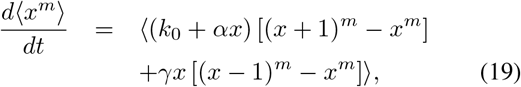

for *m* ∈ {1, 2, … } [39]–[41]. Setting *m* = 1, the dynamics of the average copy number ⟨*x*⟩ is described by the following first-order system.

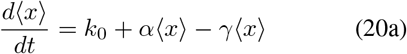

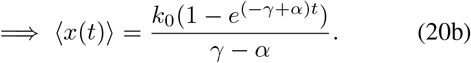

In the limit of small fluctuations, the mean FPT ⟨*T*⟩ can be approximated by the time t at which ⟨*x*(*t*)⟩ = *X* yielding

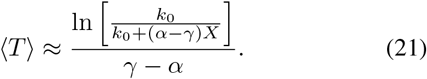

Considering this linear feedback, while keeping ⟨*T*⟩ constant, we aim to find the optimal α that minimizes the noise in *T*. To fix ⟨*T*⟩ in the low-noise limit, we adjust *k*_0_ according to *α*, from (21), as follows.

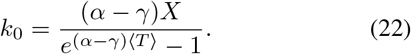

The variance in molecular counts is obtained analogously using *m* = 2 in (19),

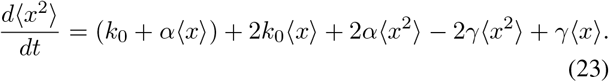

In this low-noise regime, the variance in FPT can be approximated as [42]

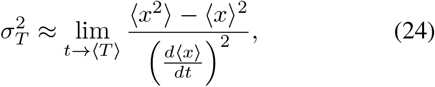

and is inversely related to the slope at which the mean trajectory ⟨*x*⟩ approaches the threshold [42]. Hence, when this slope is *flatter*, the noise of threshold-hitting times is amplified. Plugging the solutions of (23) into (24) and using (20a)-(22), we obtain the following analytical expression for the coefficient of variation of *T*.

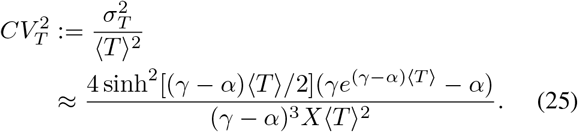

In the absence of degradation and assuming to have no feedback (*γ* → 0 and *α* → 0), the formulas (21) and (25) reduce to the following exact forms

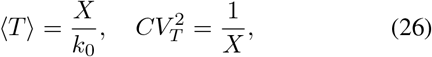

respectively, which are consistent with (10) and (11).

Fig. 3 illustrates the effect of feedback strength *α* on 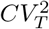, while keeping ⟨*T*⟩ constant by adjusting *k*_0_ according to (22). The analytical approximation of 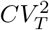 given by (25) matches well with the exact values from stochastic simulations, especially at low noise levels. As predicted, the discrepancy increases with higher noise. Moreover, 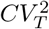 is a concave function of *α* with a global minimum. The optimal *α*, denoted by *α*^*^ which minimizes 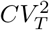 is obtained by solving the equation:

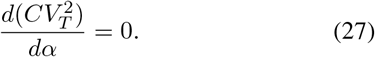

Our analysis approximated the *α*^*^ as

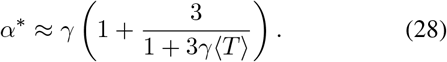

**Fig. 2:**
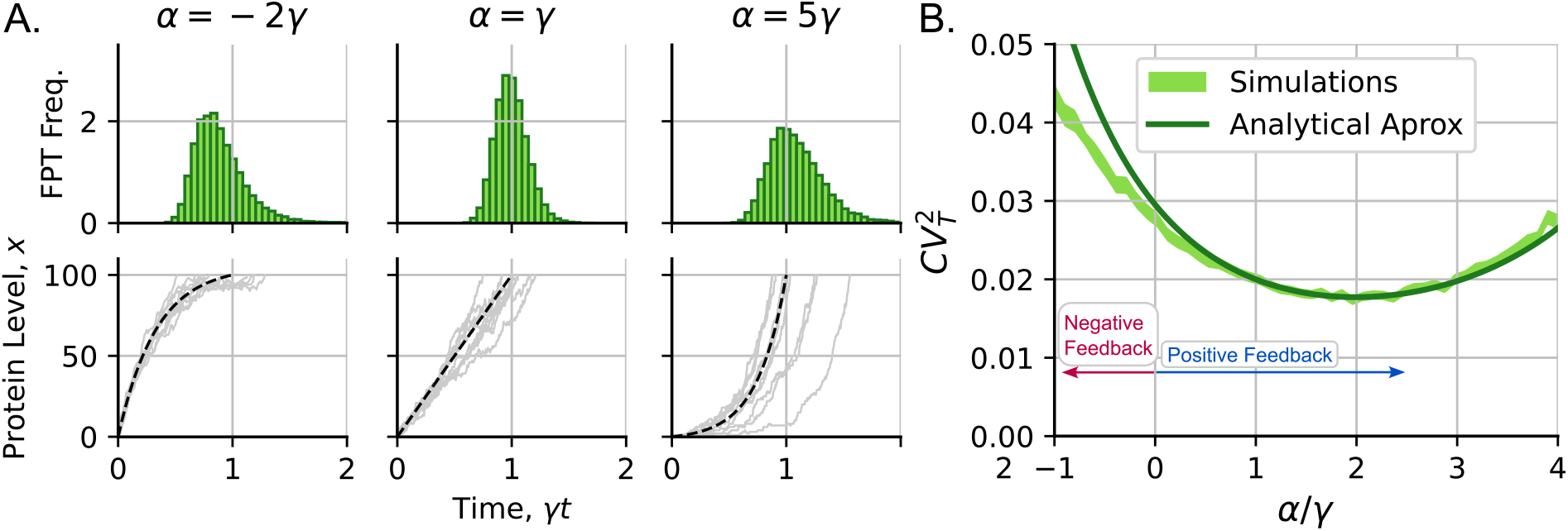
Stochasticity in FPT for linear protein synthesis rates and spontaneous protein degradation with rate *γ*. The noise of FPT is minimized using an optimal feedback strength *α*. **A**. Protein accumulation for different feedback strategies *left*: *α* = − 2*γ, center*: *α* = *γ, right*: *α* = 5*γ. Top*: FPT histogram. *Bottom*: Some illustrative trajectories of protein level *x* as a function of time. **B**. The noise in FPT (measured by its coefficient of variation *CV*_*T*_ is plotted as a function of *α*. The analytical expression (25) is compared with *CV*_*T*_ values obtained by performing stochastic simulations (the line width shows the 95% confidence interval from 10,000 simulation replicas). The parameter values used for this graph are *γ* = 1, *k*_0_ such that ⟨*T*⟩ ≈ 1, *x*(0) =0, and *X* = 100.

**Fig. 3:**
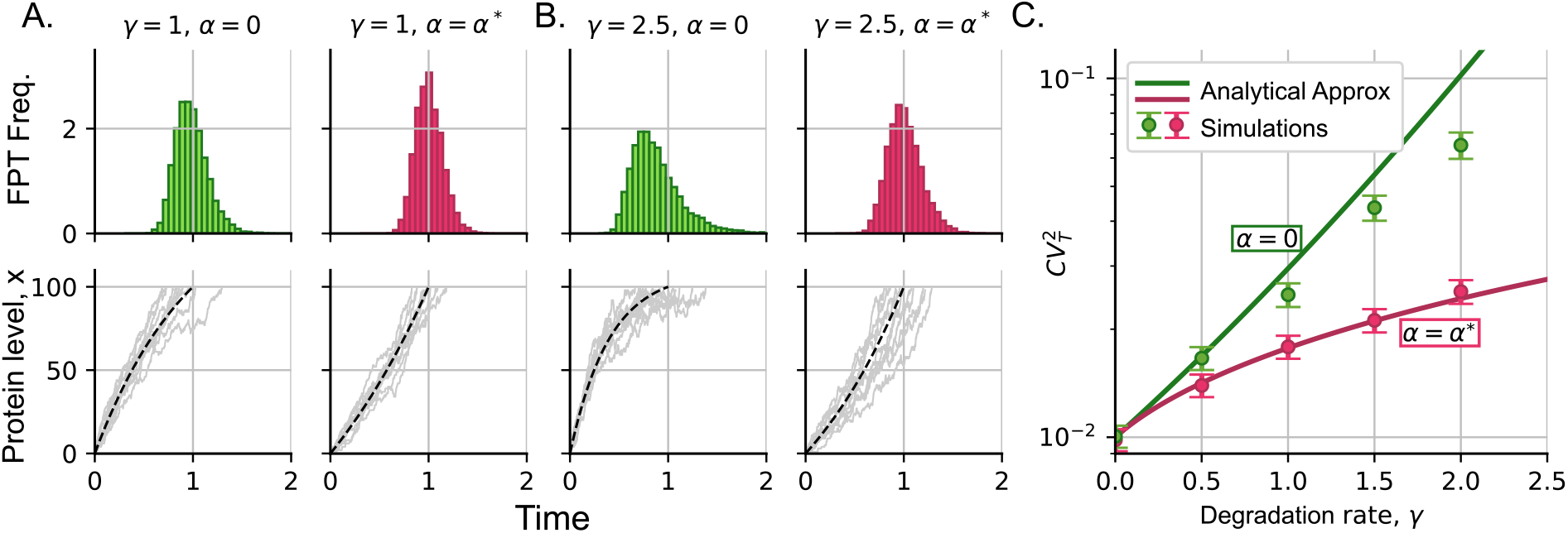
Stochastic variability in the FPTs monotonically increases with species degradation rate. **A**. Illustrative examples of accumulation processes changing with degradation rate *γ* and the feedback strength *α* maintaining ⟨*T*⟩= 1. *Top*: FPT histogram. *Bottom*: Some illustrative trajectories of the protein level *x* as a function of time. **B**. Same as A. but with *γ* = 2.5. **C**. Noise in FPT measured by the squared coefficient of variation 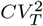 versus *γ* keeping ⟨*T*⟩ fixed. Green represents the noise without feedback (*α* = 0) and red represents the optimal linear feedback *α* = *α*^*\**^ defined in (28)). The noise is calculated from the analytical expression (25), *x*(0) = 0, *X* = 100. Error bars represent the 95% confidence intervals of the simulation results using bootstrapping methods of 2000 replicas.

This optimal solution implies positive feedback *α*^*^ > 0 and depends on the dimensionless factor *γ* ⟨*T*⟩, which is the mean FPT normalized by the average protein lifespan 1/*γ*. Fig. 3 compares the changes in 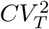 with *γ* for no feedback (*α* = 0 in green) and optimal feedback (*α* = *α*^*^ in red) with *α*^*^ as given by (28). As *γ* → 0, we see that *α*^*^ → 0 and both lines converge to 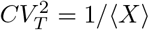, the fundamental lower bound of timing noise. For a fixed mean ⟨*T*⟩, the timing noise increases with *γ*, but more slowly for *α* = *α*^*^ (red) than for *α* = 0 (green).

## V. Optimal feedback strategy for arbitrary synthesis rates

We now relax the assumption of a linear form to allow the synthesis rate to vary without restriction. Given the analytical intractability of this problem, we focus primarily our analysis on low threshold levels and zero initial conditions to get an idea of the optimal form of *k*_*i*_. We first consider a threshold of *X* = 2 molecules where the optimization problem has an exact analytical solution.

### A. Threshold of two molecules

The simplest system that includes protein degradation is presented in Fig. 4A consisting of the synthesizing of two molecules with life-span with mean 1/*γ*. In this case, *X* = 2. Hence, we have to solve to two rates *k*_0_ and *k*_1_ that will minimize ⟨*T* ^2^⟩ for a fixed ⟨*T*⟩. To find the probability density function (pdf) of the first-passage time *T* we can write the Chemical Master Equation (CME) [43]–[46]

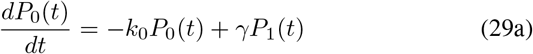

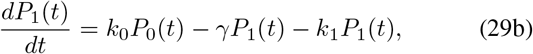

that describes the time evolution of the probabilities

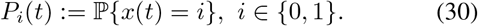

**Fig. 4:**
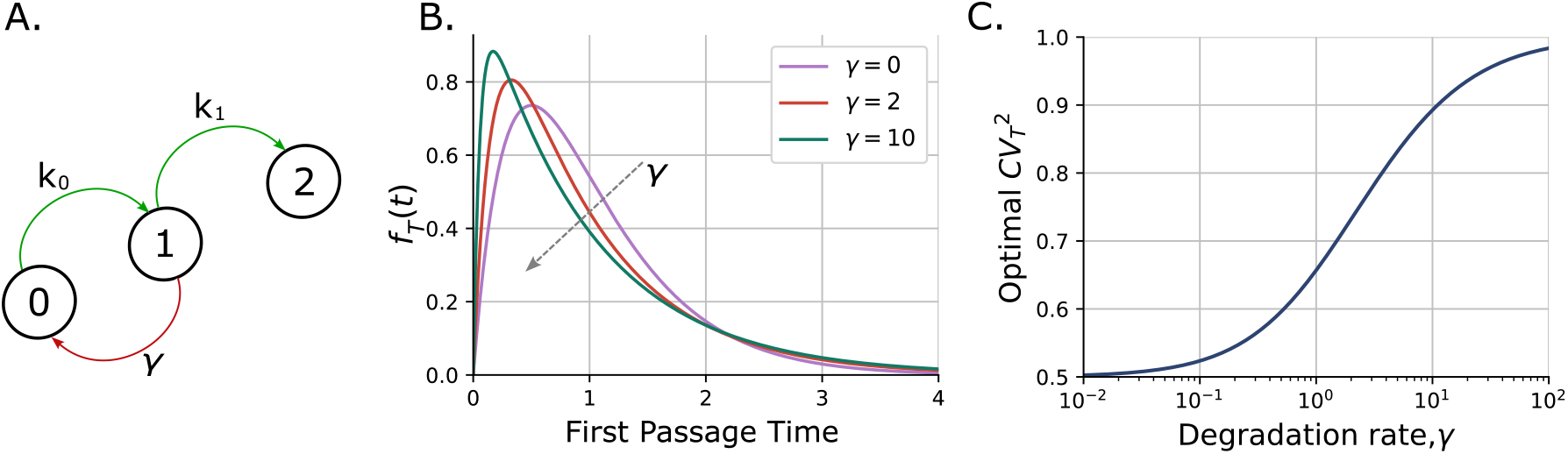
Optimal protein synthesis strategy for a threshold of two molecules considering a degradation rate *γ*. **A**. Schematic diagram where, starting from zero proteins with the objective of reaching two molecules, we optimize the rates *k*_1_ and *k*_0_ such that we minimize the variability in the FPT *T* for fixed ⟨*T*⟩. **B**. The optimized FPT distribution *f*_*T*_ (*t*) with miminial timing noise for different degradation rates *γ*. The optimal strategy corresponds to both rates being *k*_1_ and *k*_0_ equal and given by (33). **C**. Optimal timing variability as given by (34) as a function of *γ*. Here, ⟨*T*⟩ = 1 and *x*(0) = 0 with probability one.

Solving the linear dynamical system (29a) with initial conditions *P*_0_(0) = 1 and *P*_1_(0) = 0 gives the pdf of *T* which is defined as *f*_*T*_ (*t*) = *k*_1_*P*_1_(*t*). Using the obtained pdf, we find the following first- and second-order moments of *T* :

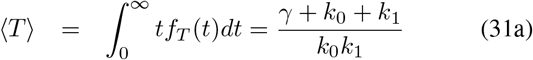

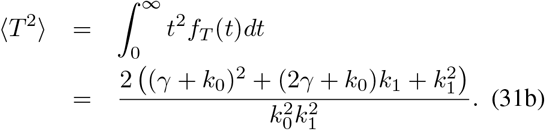

From (31a), a fixed ⟨*T*⟩ can be achieved by choosing:

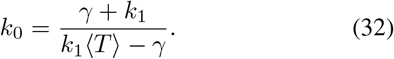

The optimization is performed on *k*_1_ that minimizes ⟨*T* ^2^⟩. The solution of this optimization reveals the optimal synthesis strategy to be

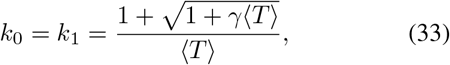

and the corresponding minimal FPT noise is

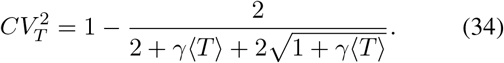

Fig. 4B illustrates the FPT distribution with rates chosen as (33) and illustrates its shape with increasing *γ*. Note that in the absence of degradation *γ* = 0, (34) reduces to 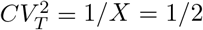. For small values of *γ* ⟨*T*⟩ ≪ 1, the noise level can be approximated by 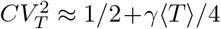. Furthermore, 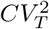 is a monotonically increasing function of *γ* ⟨*T*⟩, with 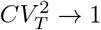 at the limit *γ*→∞ (Fig. 4C). Recall that the CV of an exponential distribution is one, and this can be seen in the shape of *f*_*T*_ (*t*) in Fig. 4B with *γ* → ∞.

### B. Threshold of five molecules

In our analysis, so far, we have found optimal strategies for some simplified cases. We derived the solution for protein accumulation without degradation, the low-noise solution for a high threshold, and the exact solution for a threshold of two molecules. However, we do not expect a simple analytical solution for the general case of protein accumulation with degradation and an arbitrary threshold. In this section, we resort to numerical methods to obtain the solution for the optimal form of *k*_*i*_ in a system with accumulation of five protein molecules, this is, *X* = 5.

To tackle this problem, we propose the general master equation:

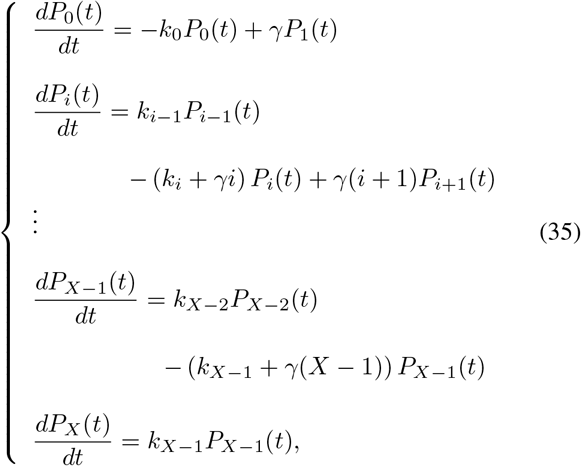

which can be used to obtain the probability vector 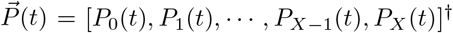 at any time *t*. Each element *P*_*x*_(*t*) represents the probability of having *x* molecules at time *t*. In a more general notation, the system can be described by the matrix **A** such as:

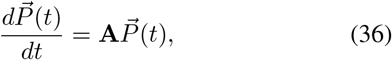

which, given the initial conditions *P*_*x*_(*t* = 0) = *δ*_*x*,0_, with *δ*_*i,j*_ being the Kronecker delta distribution, can be solved using matrix exponential methods:

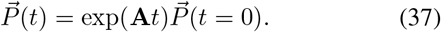

The moments of the FPT distribution can be estimated from the FPT pdf *f*_*T*_ (*t*) which, using (35), follows:

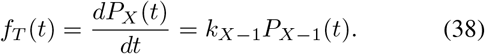

We numerically solve the moments from the FPT distribution (38) and using brute-force search we find the set of *k*_*i*_ that minimizes the 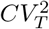 for a fixed ⟨*T*⟩ with a level of accuracy of 0.01 on both variables. We considered three cases: no feedback (constant synthesis rates); linear feedback as in Section 4; and unconstrained feedback, where any arbitrary set of *k*_*i*_ is allowed.

Fig. 5 presents and contrasts these three different cases for *γ* ∈ {0, 1, 2} with the normalized time, such that the mean FPT follows ⟨*T*⟩ = 1. In the case without feedback (Fig. 5A), the constant rate is simply set by ⟨T⟩. In the particular case of *γ* = 0, these rates are given by (10). When the feedback is constrained to follow a linear function of the protein levels, we observe that the optimal slope is positive (positive feedback). Moreover, the slope of this relationship increases with *γ*, which is consistent with our earlier approximate analytical analysis. In Fig. 5C, where the rates are unconstrained, the optimal feedback involves first increasing the rates with protein numbers. These rates reach a maximum just before the threshold. Finally, the synthesis of the last molecule to reach the threshold can be done at a much lower rate. This implies that the optimal strategy here is mixed, positive feedback at the beginning and negative feedback close to threshold. A comparison of the three cases in Fig. 4D shows the additional noise suppression achieved as we relax the constraints on the synthesis rate.

**Fig. 5:**
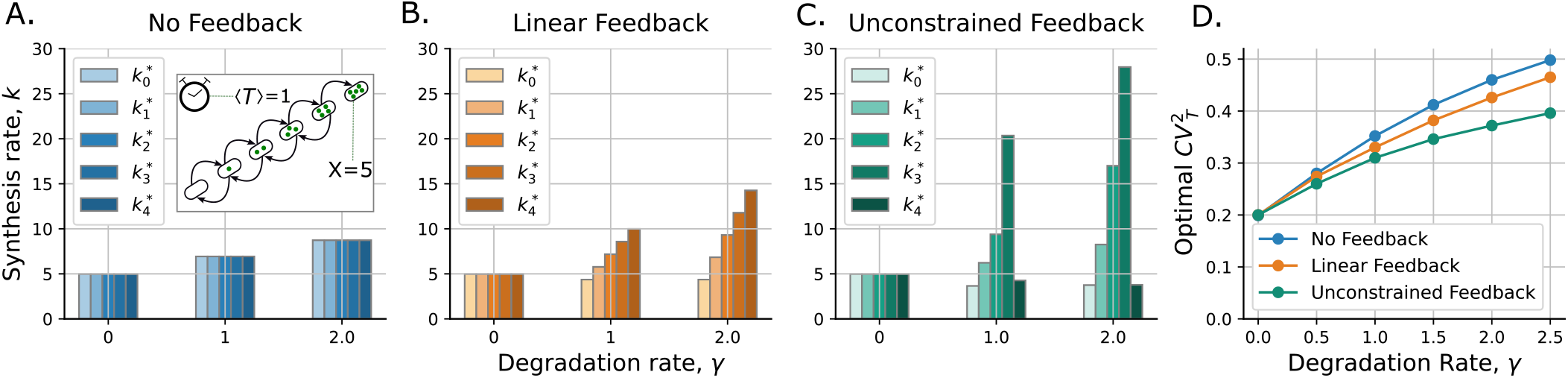
A mixture of positive and negative feedback leads to the lowest optimal noise in FPT when protein degradation is considered. Optimal synthesis rates 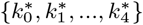 for the threshold of five molecules starting with no molecules. **A**. Optimal rates considering a constant rate for all protein levels. (Inset: a diagram explaining the process of accumulation and degradation). **B**. In the case of linear feedback, the optimal synthesis rate increases with the protein level, consistent with our low noise limit analysis in Section IV. **C**.Considering unconstrained feedback, where any arbitrary set of *k*_*i*_ is allowed, results in a nonmonotonic form with first increasing synthesis rates, and then sudden decrease just before threshold. In the absence of degradation (*γ* = 0), the optimal synthesis rate is *k*_*i*_ = *X*/ ⟨*T*⟩ regardless of the feedback strategy considered, as explained in Section III-A. **D**. Increasing the degradation rate *γ*, the accuracy of the timer decreases since the noise 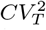 in timing increases. A comparison of optimal noise in FPT for the three strategies. For this plot, ⟨*T*⟩ = 1 and *x*(0) = 0 with probability one.

## IV. Conclusions

Accumulation of gene product levels is a key mechanism of biological timekeeping underlying diverse processes. For example, bacteriophages (viruses of bacterial cells) make proteins such as holin, that slowly build up in the host cell membrane, and the cell is lysed open upon holin’s attaining a threshold level [47]–[49]. Experimental evidence points to viruses lysing cells around an optimal time to maximize fitness [50], thus motivating the problem we have considered. Timing of cell-cycle events, such as start of DNA replication and mitotic division into daughters is also strongly influenced by species levels reaching thresholds [51]–[57].

We reviewed previous results that when proteins are long-lived, the optimal accumulation strategy is to have a constant synthesis rate that minimizes noise in timing for a given fixed mean FPT [27]. In contrast, random initial conditions shift the strategy to negative feedback regulation with synthesis rates decreasing with increasing molecule counts. With protein degradation and forcing the feedback synthesis rate to have a linear form, we show that the optimal slope of the synthesis rate is positive and monotonically increases with *γ* (Fig 2). As one would expect, the accuracy of timing deteriorates with shortening protein half-lives, and in the limit of a highly unstable protein one cannot do better than timing approaching an exponentially-distributed random variable with *CV*_*T*_ →1 (Fig. 3).

Finally, we study numerically the global optimal synthesis strategy for a five-molecule accumulation. With increasing the degradation rate, we found a pattern in the dependence of optimal synthesis rates on protein levels. The optimal strategy consists of increasing the rate with protein levels until the last molecule synthesis. The accumulation rate of the final molecule should occur at a relatively low rate (Fig. 5). As part of the future work, we plan to use some of our intuition gained for small thresholds to build an analytical theory for any arbitrary threshold. Future work on this topic could also explore other sources of variability, such as noise from cell-cycle dependent factors [58], [59]. Moreover, it would be interesting to investigate how protein interactions with *decoys*, which can buffer protein copy number fluctuations [60], [61], affect timing variations.

